# Rapid identification of microbial pathogens and antimicrobial resistance from bloodstream infections using long-read sequencing

**DOI:** 10.64898/2025.12.12.694010

**Authors:** Nicole Lerminiaux, Ken Fakharuddin, Heather J Adam, Amrita Bharat, George R Golding, Irene Martin, Michael Mulvey, Laura Mataseje

## Abstract

The gold standard for bloodstream infection (BSI) diagnostics involves culturing positive blood cultures (BC) using phenotypic methods for organism identification and antimicrobial resistance (AMR) testing, which can take up to five days. However, it is crucial to optimize antimicrobial therapy as soon as possible to reduce morbidity and mortality. We present a novel laboratory and bioinformatic workflow to rapidly identify bacterial and fungal organisms and AMR determinants from positive BCs using Oxford Nanopore Technologies long-read sequencing. Using a robust clinical sample size (n=307), after a BC has flagged positive, our average turnaround time from DNA extraction to determination of species identity was 4.4 h for a multiplex run of 12 BCs, and 3.7 h for a single sample run. We demonstrated that our pipeline taxonomic species identification results agreed with conventional MALDI-TOF identification for almost all positive BCs (97.7%, 300/307). Most species were accurately identified within the first hour of sequencing (93.7 %, 281/300). We explored AMR detection for clinically relevant antimicrobials and observed that assembly-based tools had higher agreement to conventional AST (81.2% after 1 h of sequencing, 89.6% after 5 h of sequencing) than read-based tools. Finally, we developed a publicly available analysis pipeline (venae) that generates a clinician-friendly HTML report, is quick to run, and can dynamically update as more sequencing data is acquired. This study demonstrates how applying rapid, real-time genomics to BSI diagnostics can support clinical decision-making and improve patient outcomes by reducing turnaround times.

**IMPACT STATMENT:** Early pathogen detection and administration of appropriate antimicrobial therapy for BSIs has major impacts on patient survival; early administration of effective antimicrobials reduces mortality, morbidity, length of hospital stay, and development of antimicrobial resistance. Rapid real-time genomics has high potential to improve clinical decision-making and patient outcomes by reducing turnaround times (TATs) while providing high-resolution data for organism identification, AMR determination and pathogen typing. Here, we present a laboratory and bioinformatic workflow that accurately identifies species and AMR determinants in positive blood cultures within several hours, which is quicker than conventional methods which can take days. This workflow is a step forward on the path towards point-of-care diagnostics and applying real-time genomics to characterize microbial infections in clinical settings.

**DATA SUMMARY:** Illumina sequencing data for matching pure isolates were deposited in National Centre for Biotechnology Information Sequence Read Archive (NCBI SRA) BioProject PRJNA1380445. Bioinformatic analysis pipeline is available here: https://github.com/phac-nml/venae.

## INTRODUCTION

Bloodstream infections (BSIs) pose a serious threat to human life due to significant incidence and mortality rates (1–3). These infections can lead to sepsis and septic shock, which are major healthcare problems that impact millions of people globally. A global meta-analysis estimated mortality rates of 26.7 % for hospital-treated patients and 41.9 % for intensive care unit-treated patients with sepsis (4). It is crucial to provide optimal antimicrobial therapy for sepsis as soon as possible, ideally within the first hour, to reduce morbidity and mortality (5,6). The 30-day mortality risk increased significantly after 12 h of inappropriate antimicrobial treatment and was further elevated with longer time periods (7). Consequently, rapid detection and identification is required to optimize antimicrobial treatment and to limit negative effects such as toxicity and selection of antimicrobial resistance (5,8,9). Furthermore, sepsis was the greatest healthcare expenditure in the U.S.A. at $24 billion USD in 2013 (10). Each BSI episode is associated with costs up to $20,000 USD in addition to general costs of hospitalization (11). Rapid detection of BSIs can reduce healthcare expenditures (9).

Blood cultures (BCs) remain the reference standard for diagnosis of BSIs. They are highly sensitive and easy to perform but suffer from limitations such as long turnaround times (TAT), risk of contamination, limited sensitivity in cases with previous exposure to antibiotics, and inability to identify some fastidious organisms (8,12,13). BCs typically flag positive in automated systems after 10-20 h of incubation (14) and it can take an additional 36 h to identify bacterial species and three days to obtain antimicrobial resistance (AMR) results using conventional methods (15–17). Several commercial test systems are available to identify AMR genes and/or causative agents of BSIs directly from positive BCs, but they typically cover only a limited panel of organisms or genes (12,13), can have high unit cost prices (12), and their TATs can range from 6 – 24 h after BCs flag positive (17,18).

The need for more rapid diagnostic strategies for BSIs to allow early start or change to appropriate antimicrobial therapy is evident. Next-generation sequencing is being explored as a clinical diagnostic tool as it can overcome some of the limitations of conventional methods by providing high-resolution, unbiased information in real time (1,13,19–21). Along with taxonomic identity and rapid TATs, whole genome and metagenomic sequencing provides comprehensive information for AMR resistance and virulence markers that is not constrained by panels (1,13). For some Gram-positive organisms, identification by genotypic methods may be enough to guide antimicrobial therapy (17), however organism identity alone for Gram-negatives without antimicrobial susceptibility testing is less useful as their resistance mechanisms can be complex (12). Case studies are emerging that illustrate how genomics can surpass clinical practice and inform clinical management for complex bacterial infections (22), as well as how routine metagenomics can be implemented to diagnose infections in intensive care units (23,24). However, barriers exist to implementing these types of technology routinely, including high costs, infrastructure requirements, complex data interpretation, accuracy of AMR gene-phenotype keys, and the need for standardized protocols (13,20,21).

Multiple recent studies have investigated the use of Oxford Nanopore Technologies (ONT) long-read sequencing to identify pathogens and associated AMR gene markers in positive BCs (25–38). These studies demonstrated TATs between 2.6 – 9 h, which is a significant improvement compared to multiple days for conventional workflows (25–30,32,35–38). Results so far have been very promising; ONT sequencing had much higher sensitivity and specificity compared to conventional culture methods (28,30,31,36). ONT sequencing can determine taxonomic identity as quick as 10 mins after sequencing starts (37), and it can better identify fastidious/uncultivable organisms and polymicrobial infections (1,30). However, previous studies often had small sample sizes, used older ONT technology, typically excluded fungal pathogens, and have not published bioinformatic workflows. Developing clear reporting formats to share these results with clinicians and microbiologists is also an important consideration for integration of these workflows into clinical settings, and reporting strategies have not been addressed in other studies.

To build on previous work, we developed a workflow to rapidly and accurately determine species identity and predict antimicrobial resistance (AMR) phenotypes from positive blood cultures (BCs). We optimized DNA isolation directly from positive BCs, sequenced DNA using Oxford Nanopore Technologies platforms, and designed a bioinformatics pipeline to identify bacterial and fungal organisms and associated AMR. Our study is unique in that we prioritized a large clinical sample collection (3-10-fold more than most previous studies), we used the newest ONT technology (R10.4.1/Dorado), we included fungal pathogens in our analysis, we released a public bioinformatic pipeline called venae (https://github.com/phac-nml/venae), and we designed a clinician-friendly HTML results report. This work demonstrates that it is possible to reduce BSI diagnostic turnaround time from days to hours.

## METHODS

### Ethics statement

This study was approved by the Public Health Agency of Canada’s Research Ethics Board (REB 2020-023P). No personal information from the patients was collected for this study. Human reads obtained from ONT sequencing were discarded and not used for any bioinformatic analyses beyond quantifying the amount (number and bases) of host reads generated.

### Blood culture samples

Blood specimens were collected in aerobic, anaerobic, and pediatric blood culture (BC) bottles and were incubated at 35 – 37 °C in a BACT/ALERT 3D automated microbial detection system (BioMérieux, Saint-Laurent, Canada) until samples flagged positive. The clinical site has an institutional policy permitting additional testing of discarded patient specimens. Each shipment contained 1 – 1.5 mL aliquots of each BC sample that flagged positive over a 24 h period. Average number of samples per shipment was 8, with a range of 4 – 12. Previous work showed a decrease in viability of BCs with increased storage times (39). Accordingly, all BC samples were extracted within 4 h of receipt to the laboratory, making a total maximum time of 28 h from flagging positive to extraction.

In total, there were four negative controls. DNA extraction from these BCs yielded minimal reads (n < 25 reads > 2 kb) containing microbial DNA (with >93.8 % of bases mapping to *Homo sapiens*).

### Conventional workflow for species identification and AST of matching pure isolates

All genomics-based species identification and AST results were compared to conventional phenotypic methods validated for clinical use at the clinical site. Species identification was determined at the clinical site by MALDI-TOF (Bruker BioTyper) on corresponding pure cultured isolates. Matching pure isolates were tested for antimicrobial susceptibilities by broth microdilution using the Sensititre Complete Automated AST System (Thermo Fisher Oxoid, USA) and minimum inhibitory concentrations were interpreted with CLSI breakpoints (40,41). The following panels were used (Thermo Fisher Oxoid, USA): NARMS (Gram negatives), GPALL1F (Gram positives), STP6F (*Streptococcus* isolates), and custom panels YCML1FCFA/YCML2FCFA (fungal isolates).

### DNA extraction and purification

We tested several extraction kits: QIAamp BiOstic Bacteremia DNA kit (QIAGEN, Germany), MolYsis Complete, MolYsis Basic + Epicentre MasterPure^TM^ Complete kits (Mandel Scientific, Canada), MOlYsis Basic + DNeasy Ultra Clean (QIAGEN), and QIAamp DNA Microbiome prep (QIAGEN). Given variable outcomes, the QIAamp BiOstic Bacteremia DNA kit (hereafter BiOstic) performed the most reliably in generating samples that could be sequenced using Oxford Nanopore Technologies (ONT) flow cells (see Results section Workflow Development). We input the total amount (1 – 1.5 mL) of BC sample received, which was slightly less than the 1.8 mL recommended by the manufacturer’s instructions. DNA was quantified with the Qubit fluorometer (Thermo Fisher Scientific, USA), and DNA purity was measured with NanoDrop (Thermo Fisher Scientific). If sample A260/A280 ratios were below 1.8 or if A230/A260 ratios were below 2.2, an additional AMPure XP bead (Beckman Coulter Inc., USA) clean up step was added which consisted of adding AMPure XP beads at a 1.8X ratio to the DNA and following the PCR Purification steps to incubate, wash and elute described in the AMPure XP guide.

For matching pure isolates, the Mag-Bind® Universal Pathogen 96 DNA kit (Omega Bio-Tek, USA) was used to extract DNA.

### Genome sequencing

Libraries for long-read sequencing of positive BCs were prepared using the Rapid Sequencing Kit (SQK-RAD114 kit, Flongle) or the Rapid Barcoding Kit (SQK-RBK114-24, MinION Mk1B) according to manufacturer’s instructions (Oxford Nanopore Technologies, United Kingdom). Samples were sequenced individually or multiplexed on R10.4.1 flow cells on either Flongle or MinION Mk1B devices. Basecalling was conducted using Dorado v0.9.0 with the super accurate model v5.0.0. Libraries were run for 24 h (Flongle) or up to 72 h (MinION Mk1B). We conducted MinION runs with adaptive sampling via depletion of host genome through MinKNOW v25.03.9, using the GRCh38.p14 human reference genome as input (accession GCA_000001405.29). Sample DNA was normalized to 200 ng or 75 ng input for MinION and Flongle runs respectively. In total, we conducted 38 Flongle runs, 29 regular runs, and 11 adaptive sampling runs. Flongle and adaptive runs were done in parallel with regular runs. Samples were multiplexed up to 24 samples per MinION flow cell (min: 3, median: 12), with a median yield of 6.3 Gbp per flow cell. Only single samples were run on Flongle flow cells, with a median yield of 42.7 Mbp.

Matching pure isolates were sequenced by preparing short-read Illumina libraries with the NexteraXT library preparation kit (Illumina, USA). Library pools were sequenced on an Illumina NextSeq 2000 platform with the 300 cycle output kits.

### Time point analysis

A time-point analysis was performed to compare sequencing data available after eight timepoints for both organism ID and AMR gene detection. Output .pod5 files from ONT sequencing were subsampled using a custom script to output reads from the first 1h, 2h, 3h, 4h, 5h, 8h, 12h, and 20h of sequencing. Accordingly, eight read subsets were obtained for each processed BC and ran through the analysis pipeline.

### Bioinformatic pipeline – read processing and assemblies

We designed a Nextflow-based bioinformatics pipeline called venae (https://github.com/phac-nml/venae), which takes reads output by the sequencer, performs analysis, and generates a clinical-friendly report. Host reads are removed and discarded with NoHuman v0.3.0 (42) using the Kraken2 Human Pangenome Reference Consortium database v1 (2023–09). Reads are then processed with NanoQ (43) to retain reads > 1 kb for assembly and reads > 2 kb for species identification (44,45). Read quality was assessed with NanoQ and NanoPlot (46). Long-read assemblies are generated with Flye v2.9.4 (47) using reads > 1 kb in length. Assembly completeness is analyzed with CheckM2 v1.0.2 (48) using database v2 (https://zenodo.org/records/5571251) and assembly coverage is calculated with CoverM v0.7.0 (49).

### Bioinformatic pipeline – organism identification

Species identification is performed with Kraken2 (50) using the PlusPF 8 Gb database (2024-12-18, including human, bacteria, protists, and fungi) and with sylph v0.8.0 (51) using the bacteria dbv1 GTDB r220 (113,104 genomes 2024-04-24) database and the fungal RefSeq v0.3 -c200 2024-07-25 (595 genomes). The “sequence_abundance” value output by sylph was used for reporting purposes. Only species with > 2% of classified reads and a minimum of 10 reads are considered and these cutoffs are customizable. The minimum of 10 reads was determined by examining read identity and count in negative controls.

### Bioinformatic pipeline – AMR prediction

Identification of antimicrobial resistance genes from reads is conducted with KmerResistance (52,53) (k-mer based) using the ResFinder database (54). Long-read assemblies are generated with Flye v2.9.4 using reads > 1 kb in length and are used to screen for antimicrobial resistance genes using StarAMR v0.11.0 (55) (includes MLST) using a minimum > 90% identity to the reference gene. ChroQueTas v1.0.0 (56) is used with default settings to query the FungAMR v1 database (57) for mutations in BCs containing fungal organisms.

### Bioinformatic pipeline – species-specific typing

BCs containing *Staphylococcus aureus* were screened for toxin genes (*eta, etb, tsst-1, sea, seb, sec, sed, see, seh, selk, sell, selq, lukK* and *lukS*) using VFDB (58) via ABRicate v1.0.1 (59) using database 2024-10-29 with a minimum > 80% identity to the reference gene. BCs containing *Streptococcus pyogenes* were screened for the M protein gene (*emm*) with emmtyper v0.2.0 (60) using the reference U.S. Centers for Disease Control and Prevention trimmed *emm* subtype database 2024-10-29 (2570 sequences) with a minimum > 95% identity to the reference gene. The modular nature of the pipeline will permit the incorporation of other species-specific tools in the future.

### Bioinformatic pipeline – report generation

The final report was generated via R Markdown and was designed to be clinician-friendly by including a main summary table at the top of the report and detailed results below. The summary table included five columns for sample number, identified organism, polymicrobial (yes/no), a QC parameter (pass/fail), and the predicted resistance phenotypes. Polymicrobial was defined as two or more species detected in one BC. The QC parameter was marked “pass” if the following conditions were met: (i) > 2% of reads and more than 10 reads were assigned to a single species, (ii) median read length > 500 bases, and (iii) median read quality score was > 10. If one or more of these criteria were not met, the sample was marked as “fail”. These cutoffs can be easily customized in the pipeline. The “Predicted resistance phenotype” column used data generated by StarAMR and ChroQueTas but only printed results for antimicrobials included in the Clinical and Laboratory Standards Institute document M100 ED35:2025 Table 1 (Tier 1 and Tier 2) (40) specific for each bacterial organism and antifungals with resistance confidence scores between 1 and 4 (57). The full list of predicted phenotypes and associated resistance genes is available in the detailed results section.

**Table 1.**
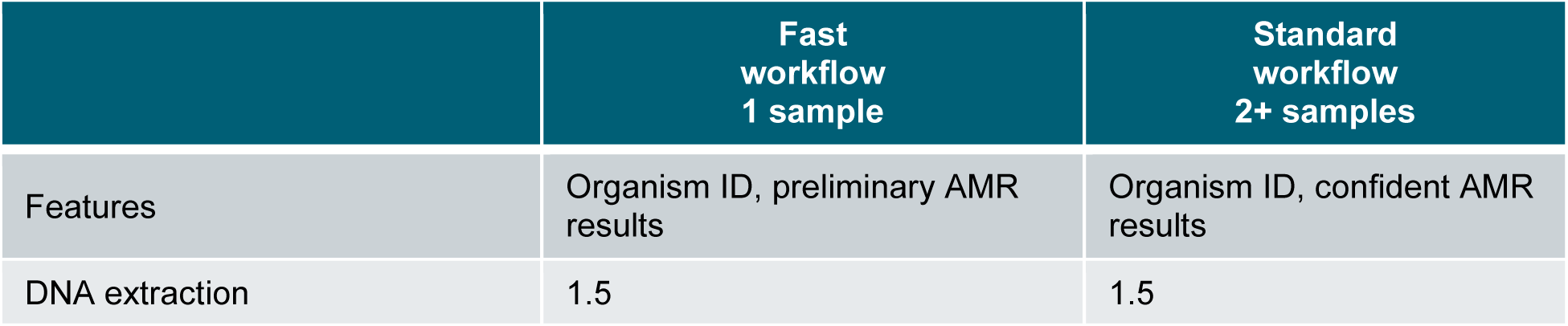
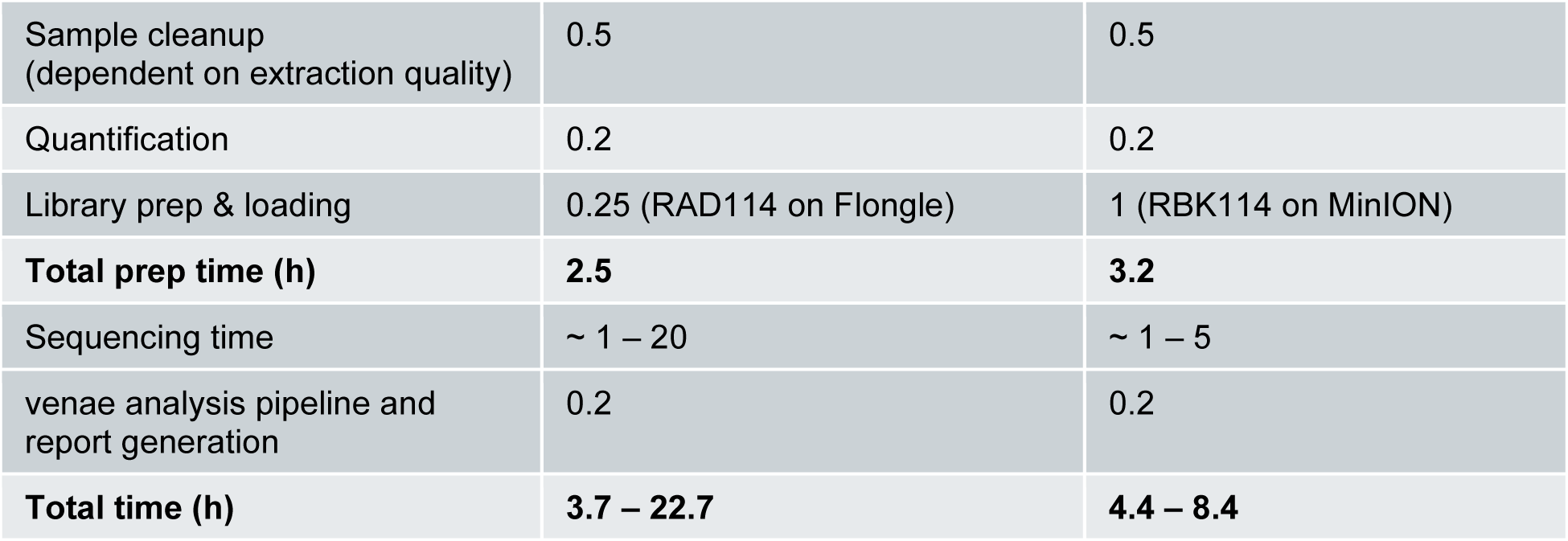
Time in hours to complete each step for three workflow options: Fast (single sample on Flongle) or Standard (2+ samples on MinION)

|  | Fast workflow<br>1 sample | Standard workflow<br>2+ samples |
| --- | --- | --- |
| Features | Organism ID, preliminary AMR results | Organism ID, confident AMR results |
| DNA extraction | 1.5 | 1.5 |
| Sample cleanup<br>(dependent on extraction quality) | 0.5 | 0.5 |
| Quantification | 0.2 | 0.2 |
| Library prep & loading | 0.25 (RAD114 on Flongle) | 1 (RBK114 on MinION) |
| **Total prep time (h)** | **2.5** | **3.2** |
| Sequencing time | ~ 1 – 20 | ~ 1 – 5 |
| venae analysis pipeline and<br>report generation | 0.2 | 0.2 |
| **Total time (h)** | **3.7 – 22.7** | **4.4 – 8.4** |

### Statistics

Accuracy was defined as (TP + TN)/(TP + TN + FN + FP), where TP (true positives) represented clinically relevant AMR phenotypes predicted by both our genomic workflow and conventional AST, TN (true negatives) represented susceptible phenotypes predicted by both the genomic workflow and by conventional AST, FN (false negatives) represented clinically-relevant AMR phenotypes detected by conventional AST that were not predicted by the genomic workflow, and FP (false positives) represent clinically-relevant AMR phenotypes detected by the genomic workflow that were not detected by conventional AST. Very major error rate was defined by FN/(FN + TP), and major error rate was defined as FP/(FP + TN).

## RESULTS

### Laboratory workflow development

Our rapid genomics workflow (referred throughout as the genomic workflow) was designed for rapid processing of positive BCs by using kit-based DNA extraction, sequencing on a rapid third-generation long-read platform (MinION, ONT) and analyzing with a dedicated pipeline. The DNA extraction and quality assessment is completed in ∼ 2 h, library preparation and flow cell loading is performed in ∼ 1 h, the libraries are then sequenced on MinION ONT platform (1 – 20 h), and sequencing output is analyzed by an analysis pipeline (∼ 20 min). Altogether, this is 3 h hands-on wet lab time, and an additional 1.5 h for organism ID and initial AMR gene predictions.

We assessed the performance of our workflow on positive BCs processed in parallel with the conventional workflow, which includes MALDI-TOF and conventional AST by broth microdilution. Matching pure isolates obtained from conventional BC processing methods were treated as references and submitted for Illumina sequencing.

We evaluated several DNA extraction kits marketed towards extracting microbial DNA directly from positive BCs. The MolYsis line of kits include a host removal step; on average, the DNA concentrations we obtained with these kits were low (median: 19.2 ng/µL) and the DNA was consistently of poor quality (median A_260_/A_230_ ratio: 1.02, median A_260_/A_280_ ratio: 1.77) (**Supplementary Fig. 1**). We attempted sequencing these extracts and experienced immediate and extensive pore death upon loading the flow cells with MolYsis libraries. Despite multiple attempts to further purify DNA following previously published protocols (26,61), we did not obtain any quality sequencing data. We also evaluated the QIAamp DNA Microbiome kit (QIAGEN) which was used with BCs previously (27) by extracting DNA according to manufacturer’s guidelines but obtained low-quality DNA (median concentration: 28.3 ng/µL, median A_260_/A_230_ ratio: 0.15, median A_260_/A_280_ ratio: 0.42) and experienced flow cell death similar to the MolYsis kits. Due to the cost of this kit, we did not pursue further troubleshooting. The BiOstic kit was the only kit that produced high-quality DNA (median concentration: 210 ng/µL, median A_260_/A_230_ ratio: 2.39, median A_260_/A_280_ ratio: 1.89) and yielded sufficient sequencing data, despite not having a specific host removal step. We tried to include an additional centrifugation step to remove host cells prior to the BiOstic kit extraction as described previously (29), but this also resulted in poor quality DNA that did not produce any sequencing data. We observed that when DNA extracted with BiOstic kits had an A_260_/A_230_ ratio below 2.2, it was beneficial to perform an AMPure cleanup on this sample to remove inhibitors that could cause extensive pore inactivation on flow cells. These cleanups did not improve sequencing performance of other DNA extracts (BiOstic DNA extracts with A_260_/A_230_ ratio above 2.2 or DNA extracts from other kits). Based on these results, we proceeded with the BiOstic kit for the remainder of this project.

### Positive BCs and host removal

A total of 307 positive BCs were processed in parallel using our genomic workflow and the clinical conventional workflow (see Methods for details). The four negative control BCs had low numbers of sequencing reads (13, 20, 12, and 29 reads longer than 2 kb respectively) and consequently no species were assigned, which agreed with conventional methods.

Our DNA isolation protocol did not include a specific host-removal step. A median of 30.3 % of reads and 36.8 % of bases sequenced from BCs were host DNA. The proportion of host:bacteria reads remained constant throughout the sequencing run.

We used a bioinformatic tool (NoHuman) to discard host reads from sequencing data. Following the recommended filtering parameter of >=2 kb read length for organism ID (44), we found that after 20 h of sequencing, the median number of remaining host reads after removal was 2 reads (median proportion 0.01 %). The median number of host reads in negative controls was 3.5. Fungal samples had the highest number of host reads across BCs, with a median number of 7.5 reads (median proportion 0.27 %).

### Taxonomic identification of BCs

Taxonomic identification from ONT sequencing agreed with conventional MALDI-TOF identification methods in the majority of BC samples (300/307, 97.7%). Most positive BCs contained a Gram-positive organism (230/307, 74.9 %) and the remainder contained a Gram-negative organism (91/307, 29.6 %) or fungal organism (11/307, 3.6 %), representing 46 Gram-positive species, 19 Gram-negatives species, and 4 fungal species (**Supplementary Table 1**). The most commonly identified organisms in BCs were *Staphylococcus epidermidis* (n=47)*, Staphylococcus aureus* (n=39)*, Escherichia coli* (n=28)*, Klebsiella pneumoniae* (n=23)*, Streptococcus pneumoniae* (n=22), and *Streptococcus pyogenes* (n=20) (**Figure 1A**). Fungal species included *Nakaseomyces glabratus* (formerly *Candida glabrata,* n=6), *Candida albicans* (n=3), *Candida dubliniensis* (n=1), and *Candida orthopsilosis* (n=1). ONT sequencing provided more accurate identification than MALDI-TOF in 60 BCs by providing species-level designations as opposed to group/complexes in some cases for *Bacillus* sp.*, Candida* sp.*, Enterobacter* sp.*, Klebsiella* sp., and *Streptococcus* sp., among others. The correct species level identity was confirmed by Illumina sequencing of the matching pure isolates and agreed 100% to ONT data.

**Figure 1.**
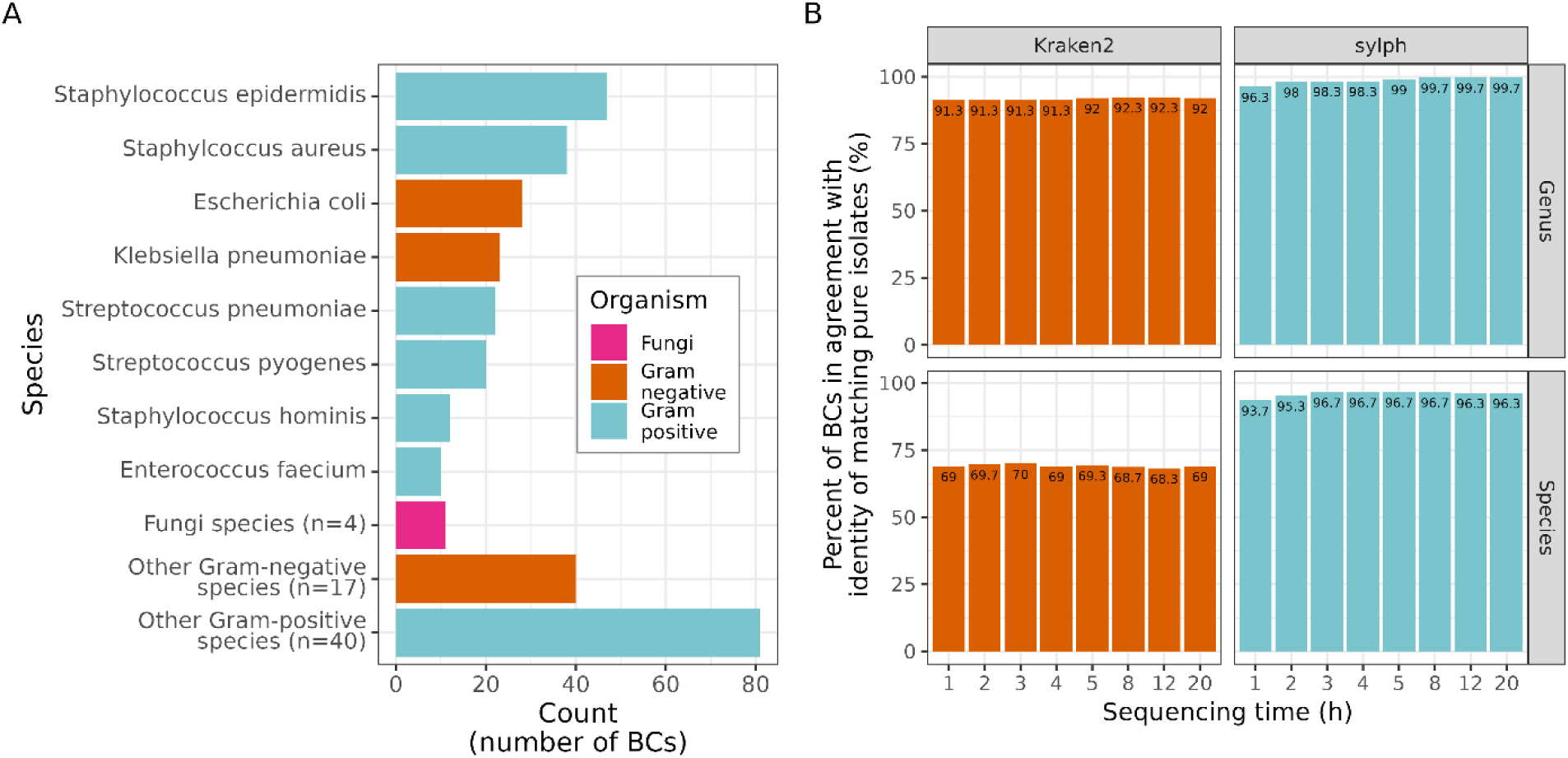
**(A)** Top species detected across 307 positive BCs. Colors indicate Gram negative, Gram positive or fungal organisms. **(B)** Proportion of BCs (n=298) in agreement with genus (top panel) and species (bottom panel) identity of matching pure isolates over time using Kraken2 (left panel) and sylph (right panel). Results from the ONT method were compared against the result of the respective tool run on the matching pure isolate Illumina sequences. BCs that did not match MALDI-TOF results were excluded (n=9). The “sequence_abundance” value from sylph was used for comparison to Kraken2.

Discordances between the genomic workflow and the conventional method (7/307, 2.3 %) consisted of identifying different species in the same genus (n=2), MALDI-TOF detecting additional species (n=3), or different genera (n=2). Two BCs had organisms that did not agree with MALDI-TOF at the species level (*Corynebacterium hessae* vs. *aurimucosum* and *Stenotrophomonas muris* vs. *maltophilia*), however others have shown that these species are not accurately identified by MALDI-TOF (62,63). The first BC with different genera identified had *E. coli* by the genomic workflow and *S. aureus* by the conventional method, and it was noted that this patient had an *E. coli* urinary tract infection two days prior to blood collection. The second BC with completely different organisms identified had *Eubacterium infirmum* (Gram-positive) by the genomic workflow and *Prevotella heprinolytica* (Gram-negative) by the conventional method. The clinical lab confirmed a Gram-positive bacilli had been observed in the initial Gram stain of the positive BC, however only *P. heprinolytica* grew.

We received 18 confirmed polymicrobial BCs from the clinical site (5.9 %, 18/307), of which the genomic workflow confirmed 15 (83.3 %, 15/18). Of the mismatches, the genomic workflow missed the second (n=2) or third (n=1) species identified by conventional methods, and in all cases, reads for the additional species were detected at 0.3 - 0.5% abundance, which is below the 2% threshold used here. Most polymicrobial cultures were a mix of Gram-positive species (55.6 %, 10/18), and the remaining were a mix of Gram-positive and Gram-negative species (44.4 %, 8/18). *Staphylococcus* sp. were frequently found in polymicrobial BCs (66.7 %, 12/18), with the most common species being *S. epidermidis* (50 %, 9/18). All but one polymicrobial BC contained two species, with the exception of one containing three (*E. coli*, *E. gallinarum, E. faecalis*). No polymicrobial cultures contained a fungal organism.

The genomic workflow identified an additional eight polymicrobial BCs that were not confirmed by conventional methods. This may be explained by the increased sensitivity of the genomic workflow, which does not rely on conventional additional culturing after the initial blood culturing for identification. Additional species present at low concentrations may produce sufficient reads for identification by sequencing.

We examined the time to species detection using our analysis pipeline venae. We used two complementary bioinformatic tools to identify organisms in BCs: Kraken2 (a classifier primarily designed for short reads) and sylph (a profiler that requires a greater read depth than Kraken2 but was more accurate in assigning species-level identity and distinguishing polymicrobial cases (51)). We treated Kraken2 as a preliminary ID and sylph as a final confident ID. Kraken2 assigned an organism identity to all BCs within the first hour of sequencing (100 %, 300/300); however, Kraken2 tended to report false positives and multiple species for a given genus. It correctly assigned genus and species for 91.3 % (274/300) and 69.0 % (207/300) of BCs after the first hour of sequencing, respectively (**Figure 1B**). This trend was stable over more sequencing time, indicating additional read data did not improve classification. Sylph requires more reads than Kraken2 to assign identity, and after the first hour of sequencing had fewer assignments than Kraken2 (96.3 %, 289/300), but a higher proportion of these assignments were correct (genus: 96.3 % (289/300), species: 93.7 % (281/300)) (**Figure 1B**). Additional sequencing time increased the percent of correctly identified BCs by sylph. Both sylph and Kraken2 suffered from database artifacts (i.e. reporting multiple species of the same genus, particularly for underrepresented organisms such as *Bacillus* and *Corynebacterium*), but these errors may be of limited clinical relevance.

### AMR predictions from sequenced BCs

We were interested in assessing the ability of ONT to correctly predict AMR from sequencing data compared to AST results of matching pure isolates. However, this dataset is biased as sample sizes for individual species are low. Consequently, this data is not a validation of AMR bioinformatic tools but rather observational. We briefly describe results for some of the major groups of organisms below, but it is important to note that these results are not comprehensive.

First, we looked at BCs sequenced with ONT vs matching pure isolates sequenced with Illumina representing all AMR genes. We excluded BCs containing fungi (n=11), BCs with mismatches in organism ID (n=5), BCs with missing matching pure isolate stocks (n=8), and BCs that did not have enough coverage for assembly (n=2), resulting in 281 BCs for AMR analysis. The proportion of correctly identified AMR genes from ONT data compared to Illumina data using assembly-based tools increased from 30.1% correct (299/992 genes representing 281 BCs) after 1 h, to 68.5 % (679/992) after 3 h, 81.9% (812/992) after 5 h, and 93.5% (928/992) after 20 h of sequencing (**Supplementary Figure 2**). This trend mirrors assembly completeness, where mean assembly completeness was 23.5% after 1 h, 62.2% after 3 h, 75.6 % after 5 h, and 91.7 % after 20 h of sequencing (**Supplementary Figure 2**). The median/mean assembly coverage values when AMR genes were missed was 6.4X/9.4X and when AMR genes were detected was 13.7X/24.1X, which indicates that 25X assembly coverage can be used as a suggested minimum coverage for accurate AMR detection.

Using read-based tools resulted in earlier detection of AMR genes, with 77.5 % correct (658/849) after 1h, 92.8 % (788/849) after 3 h, 91.3 % (801/849) after 5 h, and 93.5 % (824/849) after 20 h of sequencing. Overall, the read-based AMR tools identified 1.7X more AMR genes than the assembly-based tool after 1 h of sequencing, but the difference was smaller by 5 h (1.1X). However, the read-based tool detected more false positives (genes absent in matching pure isolates) than assembly-based tools (**Supplementary Figure 2**). While read-based tools can identify AMR genes quicker than assembly-based tools, the assembly-based tool is recommended for confident AMR prediction, as assembly requires multiple reads covering an AMR gene, which increase the likelihood that this gene is present in the BC.

We observed one instance of a BC containing a plasmid encoding antimicrobial resistance in the genomic workflow which was missing from the short-read data of matching pure isolates, indicating a likely plasmid loss. We did not observe any instances of AMR alleles not matching when average genome coverage was > 2X.

We then focused on genes with predicted AMR to antimicrobials listed in Tier 1 & Tier 2 in organism-specific Table 1 in CLSI (40). This excluded an additional 41 BCs solely comprised of organisms not represented in CLSI document M100, resulting in 240 BCs. Overall, accuracy of the genomic workflow for both assembly-based and read-based prediction tools matched the conventional AST method; read-based prediction accuracy was around 87 % at all timepoints, whereas assembly-based accuracy was 81.2% after 1 h, 87.6 % after 3 h, 89.6% after 5 h, and 91.0 % after 20 h of sequencing time (**Figure 2A**). The very major error (VME) rate (missing clinically-relevant AMR phenotypes by the genomic workflow where resistance by conventional AST was observed) decreased with more sequencing time as the major error (ME) rate (predicting clinically-relevant AMR phenotypes by the genomic workflow to which there was no resistance by conventional AST) slowly increased. We explored assembly-based results of the genomic workflow compared to conventional AST for major organism groups below. Note that the venae pipeline can produce reports in real-time throughout the run so that the end-user does not have to wait until genome completion to view results. Relevant AMR genes may be detected early in the run before genome completion.

**Figure 2.**
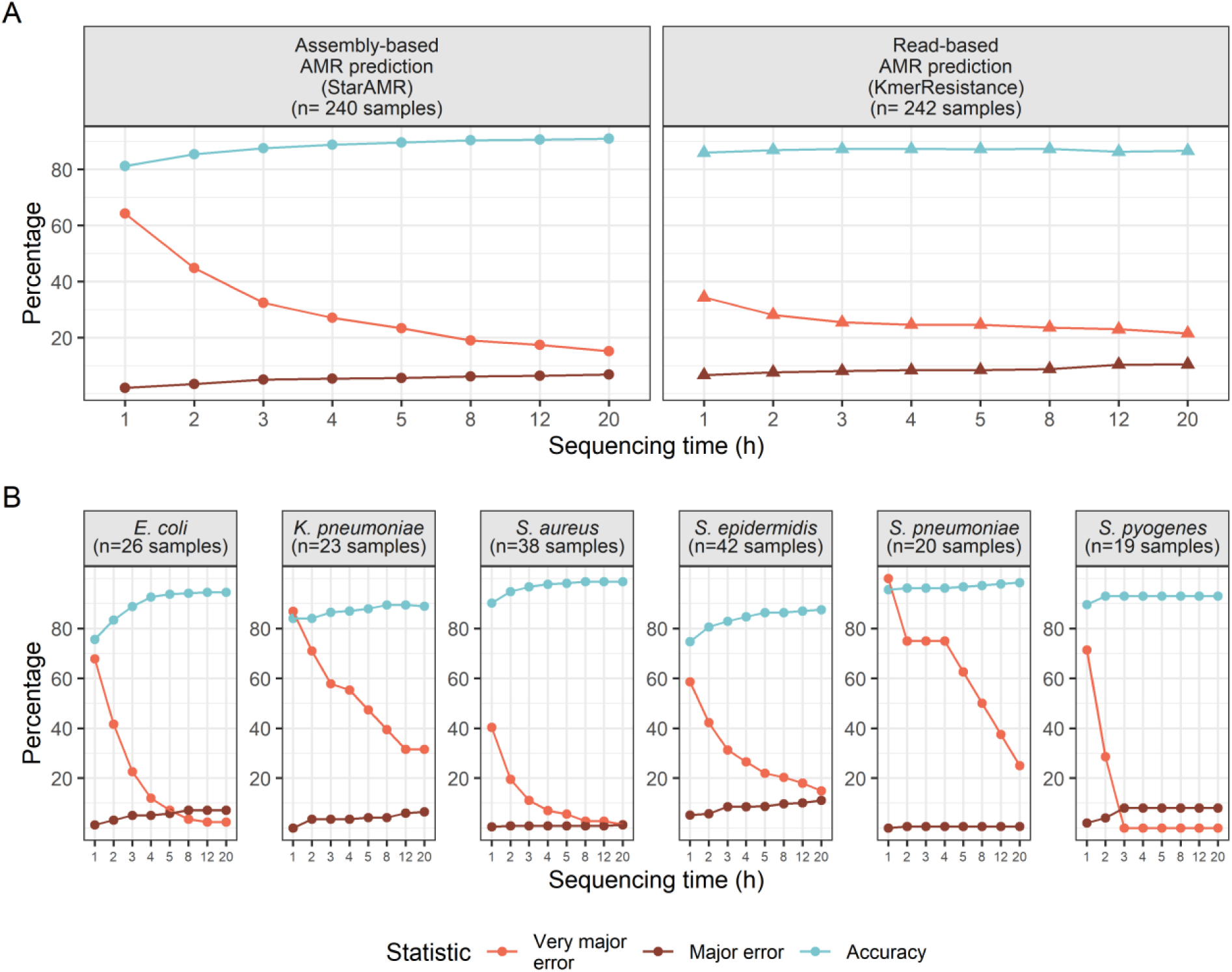
**(A)** Performance of sequencing-based AMR prediction (WGS method) using either assembly-based (left panel, StarAMR) or read-based (right panel, KmerResistance) tools compared to AST of pure culture isolates over time. BCs containing fungi (n=11) or mismatches in organism (n=5) or missing stocks (n=8) were excluded, along with BCs containing organisms not represented in CLSI M100:ED35 (n=41). Two BCs did not have enough reads for assembly-based AMR prediction. (**B**) Assembly-based AMR prediction compared to AST of pure culture isolates for top organisms. Polymicrobial BCs were excluded from this plot.

### Escherichia coli

Accuracy of predicted AMR phenotype by ONT sequencing method to conventional AST in BCs containing *E. coli* (n=26) was 75.6 % after 1 h, 88.8 % after 3 h, and 93.7 % after 5 h of sequencing time (**Figure 2B**). VME was 67.9 % after 1 h, 22.6 % after 3 h, and 7.1 % after 5 h of sequencing time. On average, it took 19.5 h of sequencing time to reach 25X coverage. Common genes identified that likely contributed to CLSI-relevant phenotypes *bla*_TEM-1B_ (ampicillin), *aac*(*3*)*-IIa* (gentamicin), *aac*(*3*)*-IId* (gentamicin), *bla*_CTX-M_ variants (ampicillin, cefoxitin, ceftriaxone), *drfA* + *sul* variants (trimethoprim-sulfamethoxazole), and *tet(A)* (tetracycline).

### Klebsiella pneumoniae

Of 23 BCs containing *K. pneumoniae*, accuracy of AMR phenotypes between the genomic workflow and conventional AST was 84.1 % after 1 h, 86.5 % after 3 h, and 87.9 % after 5 h of sequencing (**Figure 2B**). VME was 86.8 % after 1 h, 57.9 % after 3 h, and after 47.4 % after 5 h. On average, it took 15.2 h of sequencing time to reach 25X coverage. Common genes identified that likely contributed to CLSI-relevant phenotypes were *bla*_SHV_ variants (ampicillin, cefoxitin, ceftriaxone), *bla*_CTX-M-15_ (ampicillin, ceftriaxone), *bla*_NDM-7_ (ampicillin, cefoxitin, meropenem), and *drfA* + *sul* variants (trimethoprim-sulfamethoxazole). High VME errors are likely due to *bla*_SHV_ genes, which are annotated as: “unknown beta-lactam” in the ResFinder database but depending on the variant can encode resistance to multiple antimicrobials.

### Staphylococcus aureus

Accuracy of predicted AMR phenotype by the genomic workflow compared to conventional AST in BCs containing *S. aureus* (n=38) was 90.1 % after 1 h, 96.7 % after 3 h, and 98.0 % after 5 h of sequencing time (**Figure 2B**). VME was 40.3 % after 1 h, 11.1 % after 3 h, and 5.6 % after 5 h of sequencing time. On average, it took 7.8 h of sequencing time to reach 25X coverage. Common genes identified that likely contributed to clinically-relevant phenotypes were *blaZ* (penicillin)*, mecA* (oxacillin)*, msr(A)* (erythromycin)*, mph(C)* (erythromycin)*, erm(C)* (clindamycin, erythromycin)*, erm(A)* (clindamycin, erythromycin), and *tet(K)* (tetracycline).

### Staphylococcus epidermidis

*S. epidermidis* was found in 42 BCs and accuracy of predicted AMR phenotype by the genomic workflow compared to conventional AST was 74.7 % after 1h, 82.9 % after 3 h, and 86.4 % after 5 h of sequencing time (**Figure 2B**). On average, it took 6.8 h of sequencing time to reach 25X coverage. VME after 1 h of sequencing time was 58.6 %, after 3 h was 31.3 %, and after 5 h was 26.6 %. Common genes identified that likely contributed to CLSI-relevant phenotypes were *blaZ* (penicillin)*, mecA* (oxacillin)*, msr(A)* (erythromycin)*, mph(C)* (erythromycin)*, erm(C)* (clindamycin, erythromycin)*, erm(A)* (clindamycin, erythromycin), and *tet(K)* (tetracycline).

### Streptococcus pneumoniae

Of 20 BCs containing *S. pneumoniae*, only 5 (25 %) showed resistance to one or more clinically-relevant antimicrobials by conventional AST. Accuracy of predicted AMR phenotype by the genomic workflow to conventional AST was 95.6 % after 1 h, 96.1 % after 3 h, and 96.7 % after 5 h of sequencing time (**Figure 2B**). VME was 100% after 1 h, 75.0 % after 3 h, and 62.5 % after 5 h of sequencing time. On average, it took over 20 h of sequencing time to reach 25X coverage. Common genes identified that likely contributed to clinically-relevant AMR phenotypes were *erm(B)* (clindamycin, erythromycin)*, msr(D)* (erythromycin), *mef(A)* (erythromycin), and *tet(M)* (tetracycline).

### Streptococcus pyogenes

Of 19 BCs containing *S. pyogenes*, only 4 (21.1 %) showed resistance to one or more clinically-relevant antibiotics. Accuracy was 89.5 % after 1 h and 93.0 % from 2 h of sequencing time onwards (**Figure 2B**). VME was 71.4 % after 1 h, and 0 % after 3 h of sequencing time and onwards. On average, it took 7.1 h of sequencing time to reach 25X coverage. Common genes that likely contributed to CLSI-relevant phenotypes were *erm(A)* (clindamycin, erythromycin), *erm(T)* (clindamycin, erythromycin), *tet(M)* (tetracycline)*, tet(O)* (tetracycline), and *mre(A)* (erythromycin).

### Fungi

We obtained 11 positive BCs that contained a fungal organism. While most fungal organisms were identified within the first 1 h of sequencing (72.7%, 8/11), there was typically not sufficient read data to produce complete assemblies by 20 h of sequencing (most assemblies < 1 Mb in size with two exceptions of 7 Mb and 14 Mb). We queried fungal organisms against the FungAMR database to search for mutations in species-specific genes that can cause antifungal resistance. Only four mutations were identified in a single BC containing *C. albicans* (two in Cyp51 and two in Mrr2), and all had low confidence resistance scores (scored 5 – 8). However, in the matching pure isolates, there were 53 mutations detected across the 11 fungal isolates, and four of these mutations had high confidence resistance scores (scored 1 – 4): two Cyp51 mutations in two *C. albicans*, one Mrr1 mutation in one *C. albicans*, and one Pdr1 mutation in one *N. glabratus*. Only the two Mrr2 mutations identified in the *C. albicans* BC agreed with the matching pure isolates. This indicates that this workflow, while appropriate for fungal organism identification, is not optimized for complete assembly in 20 h and BCs that are identified as fungal should be sequenced longer for better resolution of AMR determinants.

Overall, species-specific AMR detection had distinct trends per organism, but generally organisms with smaller genomes and fewer AMR genes (*Staphylococcus* sp.*, Streptococcus* sp.) took less time to identify all relevant AMR genes compared to organisms with larger genomes and more AMR genes (*Enterobacterales,* fungi).

### Species-specific typing

We included two typing tools in our analysis pipeline: *emm* typing in *S. pyogenes* and toxin typing in *S. aureus*.

We detected ten different toxin genes across 38 BCs containing *S. aureus*, with 30 of these containing one or more toxin genes. Ninety percent of all toxin genes were detected correctly after 3 h of sequencing (62/69, 89.9 %) (**Supplementary Figure 3A**). The *lukF-PV* genes were the most common and were found in 73.6 % (28/38) of *S. aureus* BCs. Only three discordant results were observed in this dataset, giving overall agreement of 92.0 % (35/38). There were two BCs where additional toxin genes were detected in the matching pure isolates: one case with *lukF-PV*, and one case with *selk* and *selq*. One BC detected *selk* by the genomic workflow but this gene was absent in the matching pure isolates stock.

We detected *emm* types in 21 BCs containing *S. pyogenes* representing 14 distinct *emm* types. Most *emm* types were detected correctly after 3 h of sequencing (16/21, 76.2%) and improved after 5 h of sequencing (19/21, 95.2%). There was only one instance at one timepoint (3 h) where the *emm* type was assigned incorrectly, but this was rectified with an additional hour of sequencing data (**Supplementary Figure 3B**). One sample had low sequencing data and did not have a type assigned by 20 h (estimated coverage: 2.7X).

### Evaluating Flongle sequencing for single BCs

We investigated the capability to process a single BC with a Flongle flow cell, as the reduced library prep time for a single sample could allow data to be generated more quickly. Flongle adapters and flow cells were removed from ONT public storefronts in May 2025 but are reportedly still available. The number of active pores on the Flongle flow cells upon sequencing start varied from 14 to 93, with an average of 61, even with flow cell storage times of up to 6 months (beyond the recommended 4 week warranty period). Of the 38 Flongle runs, 8 had pores counts below the recommended warranty of 50, however there was no relationship between pore count, storage time, and amount of data output.

We selected a diversity of organisms and both monomicrobial and polymicrobial BCs for analysis, and the data output from the Flongle flow cells was variable with a median of 42.7 Mb (range: 1.22 Mbp – 169.94 Mbp). Species were correctly identified for 65.8 % (13/38) of samples by 1 hr and for 94.7 % of samples (36/38) by 4 h of sequencing time. After 20h of sequencing, assemblies were on average only 44.0 % complete (**Figure 3**). In the best performing run, assembly was 100% complete after 4 h of sequencing (*S. pyogenes*). AMR genes were detected on Flongle runs, however due to low data output, many genes remained undetected by both assembly-based and read-based tools after 20 h of sequencing time. Given the pore count of Flongles and its capacity for data generation, it is recommended for rapid identification only.

**Figure 3.**
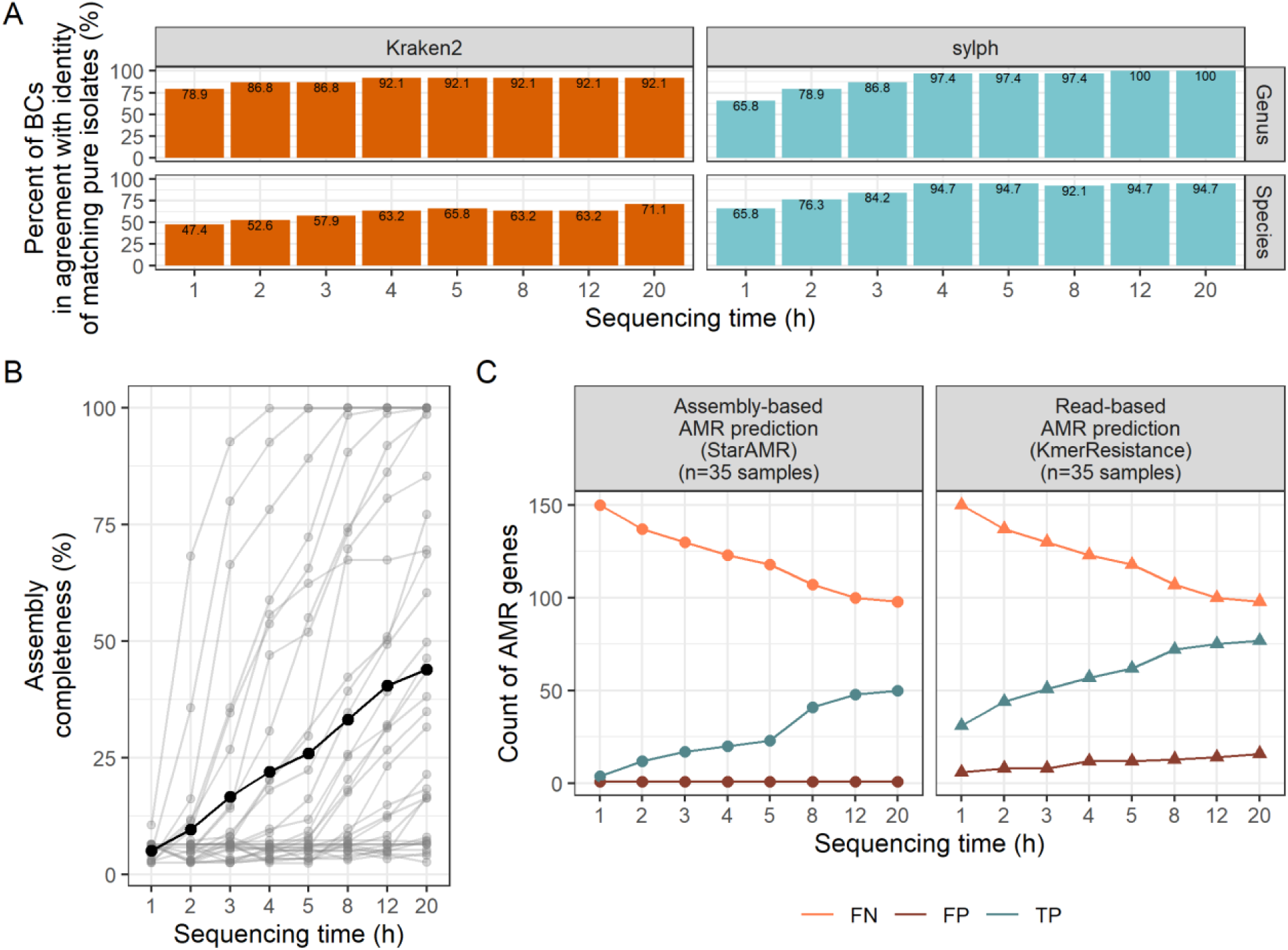
Evaluating single BC processing on Flongle flow cells (n=38 BCs). (**A**) Proportion of BCs in agreement with genus (top panel) and species (bottom panel) identity of matching pure isolates over time using Kraken2 (left panel) and sylph (right panel). (**B**) Assembly completeness over time. Bold black line indicates the mean, and grey lines represent individual samples to show variation. (**C**) Count of AMR genes detected by assembly-based tool (StarAMR, left panel) and read-based tool (KmerResistance, right panel). Three fungal BCs were excluded from this plot. The reference dataset was matching pure isolates sequenced by Illumina and assemblies analyzed for AMR genes with StarAMR and KmerResistance. FN = false negative (gene missed in ONT method), TP = true positive (gene correctly detected), FP = false positive (gene detected in ONT method that was not detected in Illumina).

### Evaluating adaptive sampling with host depletion for multiple BCs

ONT adaptive sampling provides an opportunity for faster data acquisition if there is sufficient computational capacity on the computer running the sequencing experiment. We conducted 11 adaptive sampling runs (n= 110 total BCs) to evaluate if host depletion improved time of acquisition of microbial data and AMR detection. We used the complete reference human genome for depletion (GRCh38, accession: GCA_000001405.29) and ran 8 – 12 samples per flow cell which produced a median data yield of 9.0 Gbp per flow cell (compared to 6.3 Gb median for non-adaptive runs).

During host depletion, both adaptive and non-adaptive runs had comparable averages of 30.1 % and 32.6 % reads removed respectively; however, adaptive samples had on average 30.0 % of total bases removed whereas the non-adaptive counterparts had 40.8 % of total bases removed (**Supplementary Figure 4A**). This indicates that the adaptive runs were sequencing 26.5% fewer human DNA bases than the normal counterparts. Assemblies from adaptive sampling runs were on average 5 % more complete and had an average of 124 kb more sequence assembled per genome after 1 hr of sequencing compared to non-adaptive sequencing runs (**Supplementary Figure 5A**). However, by 8 hr, adaptive sampling assemblies were 12.9% less complete and had 44 kb fewer sequence assembled compared to non-adaptive sequencing. Adaptive sampling runs detected on average only 0.24 more AMR genes after 1 hr of sequencing compared to non-adaptive sequencing (**Supplementary Figure 5B**). However, by 5 hr, adaptive sampling runs detected on average 0.5 fewer AMR genes than normal runs. Species-level identity was worse (more incorrect detections early on) for adaptive runs for both tools compared to the normal run counterparts (**Supplementary Figure 5C**). While there were variable amounts of data generated for each run in comparison with the equivalent non-adaptive run (**Supplementary Figure 4B**), the adaptive runs generally accumulated microbial data more quickly than normal runs at the beginning of the run. As time progressed, data output by the normal runs overtook any benefit obtained from adaptive sampling. As data above showed optimal AMR prediction would require more than 1 h of sequencing time, adaptive sampling would not be recommended over non-adaptive in this workflow.

### Bioinformatic workflow – venae

We developed a Nextflow-based bioinformatic workflow (named venae) that takes reads as input and outputs a final clinician-friendly HTML report (**Figure 4**). venae is publicly available on GitHub (https://github.com/phac-nml/venae) and an example HTML report can be viewed online (https://phac-nml.github.io/venae/tests/test_data/report.html). Sequencing time to obtain AMR results can vary depending on the size of organism genome and number of AMR genes present, and venae pipeline is fluid and allows for reporting of multiple organisms and samples while others are still sequencing. The pipeline requires demultiplexed ONT sequencing reads as input which can be updated as the sequencing run progresses, and the pipeline runs quickly with 12 or fewer samples (10 m runtime).

**Figure 4.**
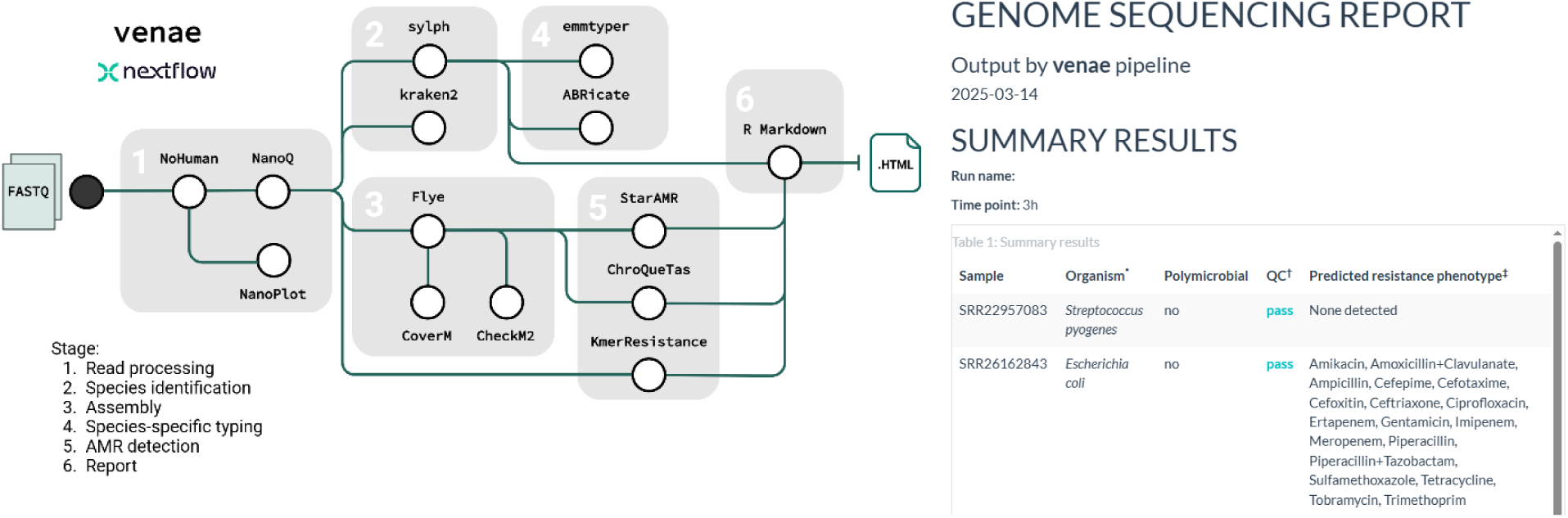
venae pipeline and output HTML report. (**A**) Overview of tools included in the pipeline. (**B**) Screenshot of Summary Table at the top of the report containing sample name, organisms identified, polymicrobial status, quality control pass/fail, and predicted AMR phenotype. Link to example HTML report here: https://phac-nml.github.io/venae/tests/test_data/report.html.

### Workflow resources (time and cost)

Our workflow had hands-on laboratory time of ∼ 3.2 h, including i) DNA extraction (1.5 h), ii) DNA quantification and purity assessment (0.2 h), iii) additional bead cleanup only if purity ratios were low (0.5 h), and iv) library prep and flow cell loading (1 h). As described above, species identity was typically confirmed within the first hour of sequencing, and AMR results were mostly confident by ∼ 5h of sequencing, although there were instances where AMR results were confirmed after 1 h of sequencing or after 12 h depending on the number of AMR genes per organism. The venae analysis pipeline, including report generation, takes ∼10 mins. Consequently, the potential turn-around time from a BC flagging positive to report generation could be achieved as early as 4.4 h when multiplexed (**Table 1**). Single-sample Flongle runs could have species identity confirmed as early as 3.7 h after flagging positive.

We calculated costs to perform both the Fast workflow (single sample) and Standard workflow (12 samples). It costs approximately $195 CAD/$141 USD to process a single BC on Flongle, and $1140 CAD/$825 USD to run 12 BCs in parallel on the Standard/MinION workflow ($95 CAD/$69 USD per sample when multiplexed) (**Supplementary Table 2**). We did not account for additional infrastructure costs including MinION sequencer/Flongle adaptor and a computer for sequencing and analysis, as well as basic lab equipment such as vortexes, thermocyclers, and heat/water baths. Note costs can vary depending on ONT website and this data is based on costs accurate at time of testing (September 2025).

## DISCUSSION

We present a novel laboratory and bioinformatic workflow to rapidly identify bacterial and fungal organisms and AMR determinants from positive blood cultures (BCs) using long-read ONT sequencing. Rapid, real-time genomics has much potential to better support clinical decision-making and improve patient outcomes by reducing turnaround times (TATs) while providing high-resolution data for organism identification, AMR determination and pathogen typing. This work provides another step towards applying real-time genomics to profile complex microbial infections in clinical settings.

Using a robust clinical sample size (307 positive BCs), our average turnaround time from DNA extraction after a BC has flagged positive to determination of species identity was 4.4 h for a multiplex run of 12 BCs. This time could be reduced further to 3.7 h by running a single sample. Our TATs fit in the range determined by other recent studies exploring ONT sequencing to rapidly diagnose positive BCs, which demonstrated TATs between 2.6 – 9 h (25–30,32,35–38). These reduced TAT represent improvements over conventional methods which take multiple days. Hong *et al.* (2023) reported that in some emergency situations a preliminary report could be produced with a 6 h TAT by using ONT sequencing if diagnostic thresholds are met, with a complete report produced within 24 h. We think this approach to early detection is valuable, and we designed our analysis pipeline to report on samples as they sequence. The pipeline will produce results for samples at various stages; high sequencing coverage for all samples is not required prior to producing a report.

This genomics workflow also has potential to support outbreak investigations. For example, different *emm* types in *S. pyogenes* isolates will rule out a link between cases immediately, and the same *emm* type can be treated as potentially related until a phylogeny can be completed. The data generated by venae can be used as inputs into other outbreak support tools (for example, WADE (https://github.com/phac-nml/wade)) to determine potential relatedness. Further development of venae could also include phylogenic typing methods such as toxin profiling or MLST of various species, though currently out of scope for this report.

Species-level identification by the genomic workflow agreed with the conventional MALDI-TOF in 97.7 % of BCs, and most were confirmed within the first hour of sequencing. This observation is similar to results from other studies (25,32,34). Kraken2 did not perform well for species-level assignment here, observed previously for simulated BCs (45) and in benchmarks with long reads (44), however we included it in our workflow as a preliminary ID tool for early reporting that is replaced by sylph when there is sufficient data for detection. We used a species identity threshold of 2 % (i.e. at least 2 % sequence abundance for a species to be identified). This threshold eliminated contaminating sequences from samples on the same run or demultiplexing errors, at the expense of three polymicrobial BCs where one confirmed species was present at 0.5 % sequence abundance. In contrast, Pilato *et al*. (2025) used a 1 % threshold, Govender *et al*. (2025) observed the best results with a 0.4% threshold, and Ali *et al*. (2024) identified 1 – 1.3 % of reads as contaminants or database artifacts. Consequently, we have made this parameter easily modifiable in the venae pipeline.

One limitation of this study was that we are only reporting on polymicrobial cases from the same BC bottle and are unable to determine polymicrobial from two separate BC bottles from the same blood draw due to our sampling strategy. Consequently, our polymicrobial BC rate of 5.9 % (18/307), which fits within the previously observed range of 3.8 – 16 % (26,27,39,45,61,64), may be an underestimate. In addition, the genomic workflow was more precise for species ID than MALDI-TOF in 60 BCs and the genomic workflow detected more species in 8 BCs that were not identified by conventional methods, consistent with other studies (28,31,36,45).

The top BC organisms identified here match the most common pathogens associated with BSIs in other studies, including *E. coli*, *S. aureus,* and other *Staphylococcus* sp. (65,66). Some BCs in this study contained organisms that can be contaminants, including coagulase-negative *Staphylococcus* species*, Streptococcus mitis, Corynebacterium* sp., *Cutibacterium* sp., *Bacillus* sp., and *Micrococcus* sp. (8,45). Every hospital will have its own guidelines on determining contaminants, but it was important for us to identify these in the workflow and generate the information to allow clinicians and microbiologists to decide how to interpret the data according to their individual guidelines. Other studies investigating BCs have focused solely on bacterial species and excluded fungi. We included fungi as part of our rapid identification and explored AMR prediction using the FungAMR database (57), recognizing the increasing prevalence of *Candida* sp. in BSIs (65). Our pipeline accurately detected fungal organisms in BCs, but not enough sequencing data was obtained to reliably detect AMR mutations in genes, and consequently conventional AST for these BCs would still be required.

AMR prediction is promising but has not yet been widely adapted into clinical use to inform patient care. An ongoing example is the implementation of respiratory metagenomics program in several centers in the United Kindown; this program has resulted in the change of antimicrobial treatment in several cases due to detection of extended-spectrum beta-lactamase resistance genes in *Enterobacterales* pathogens (24) or early institution of barrier precautions after detection of vancomycin-resistant *Enterococcus* (23). One of the greatest challenges in using genomic AMR data to support patient care is the accuracy of AMR gene-phenotype keys (67,68). Currently there are discrepancies between conventional AST and resistance predicted by genomic tools, and it is critical to have databases with reliable gene-phenotype keys to accurately inform appropriate treatment (69). AMR genes with novel/undescribed mechanisms (“unknown phenotype” in the ResFinder database or missing from the database altogether) are not captured by this workflow and are a limitation of this and all other AMR prediction analyses. For example, many *bla*_SHV_ alleles in *Klebsiella* sp. observed in this study are missing phenotypes in the ResFinder key, and certain organisms are better represented in these databases than others (32). AMR database validation is beyond the scope of this study but is essential for accurate predictions that can be actionable from a clinical perspective.

We explored read-based and assembly-based AMR prediction, although it is important to note that this dataset is biased to a single hospital site and sample sizes for individual species are low. We found that complete AMR profiles (relative to matching pure isolates) were obtained with a mean assembly coverage of 25X based on the size of reference genomes. This agrees with Weinmaier *et al*. (2023), who found that AMR results stabilized at 30X coverage, and with Liu *et al*. (2023), who calculated that the average amount of sequencing data required for accurate AMR calls was 94.6 Mb, which is approximately 19X coverage for *Enterobacterales*, 32X coverage for *Staphylococcus* sp., and 48X for *Streptococcus* sp..

Another limitation of this study was that there was no specific host depletion step, and consequently a large amount of host DNA could reduce sequencing coverage of bacterial sequences (29). We assessed several DNA extraction kits and methods published by others to remove host DNA but were unable to replicate. We found bioinformatic methods to be effective in removing host DNA (42), but evidently sequencing more microbial reads sooner would improve TATs. Recently other studies have published alternative lab-based host removal protocols such as using a combination of BiOstic and MolYsis kits (35) or a modified saponin-based protocol (33,70,71) and continuing to experiment with these host removal strategies would likely decrease the TAT of our workflow.

We proposed a workflow suited to individual samples using Flongle to overcome the barriers around current sequencing technologies that are suited to batching multiple samples to streamline throughput and drive down costs (21). As others have observed, we experienced variability in output depending on the sample and often there was not enough output for consistent AMR gene detection (35,72). Flongle runs may be best suited to organisms with smaller genome sizes (*Streptococcus* sp.*, Enterococcus* sp.*, Staphylococcus* sp.) as less output data is required for a complete assembly; however, it is impossible to know the identity of the organism ahead of time given the nature of the BCs and its more reliable to use Flongle for rapid identification purposes if that is the priority.

Despite sequencing fewer host bases, we did not observe substantial benefits using adaptive sampling with host depletion. Similar to this study, Sajib *et al*. (2024) found adaptive sampling resulted in less host DNA being sequenced but no bacterial enrichment. ONT released a new adaptive sampling guide in November 2024 with recommendations that depletion adaptive sampling will provide the most enrichment benefits when the sequence of interest is a small proportion of the sample (below 5%). Most BCs in this study were far above this threshold (average: 67.7%), likely due to the blood culturing period prior to flagging positive, which could explain why adaptive sampling likely did not provide large benefits. Adaptive sampling appears to be more suited towards sequencing samples with higher proportions of host DNA, such as directly from whole blood (73) or from respiratory samples (74).

In conclusion, we present the development and application of a laboratory and bioinformatic workflow to rapidly identify bacterial and fungal organisms in positive BCs using long-read sequencing. We demonstrated TATs for species identity as 4.4 h for multiplexed samples and 3.7 h for single samples, which significantly reduces the TAT of conventional methodologies and provides further support for applying real-time genomic tools to BSI diagnostics. We have made our analysis pipeline publicly accessible (venae) to increase accessibility and transparency and we also evaluated single sample vs multiplexing options to reduce costs. Further improvements would include continued validation of clinical AMR phenotypes from BSI isolates to improve AMR prediction tools. Testing these workflows in clinical settings, as done for respiratory pathogens (23), is the next step to inform implementation into routine clinical practice.

## ACKNOWLEDGEMENTS

This project was funded by the Public Health Agency of Canada’s Genomic Research and Development Initiative. We would like to thank Karen Wake for coordinating sample shipments and Mandy Reimer, Angela Yuen, Lisa Li, Marith Been, John Merluza, and Jonelle Tinsley for performing AST.

---

**Supplementary Figure 1.**
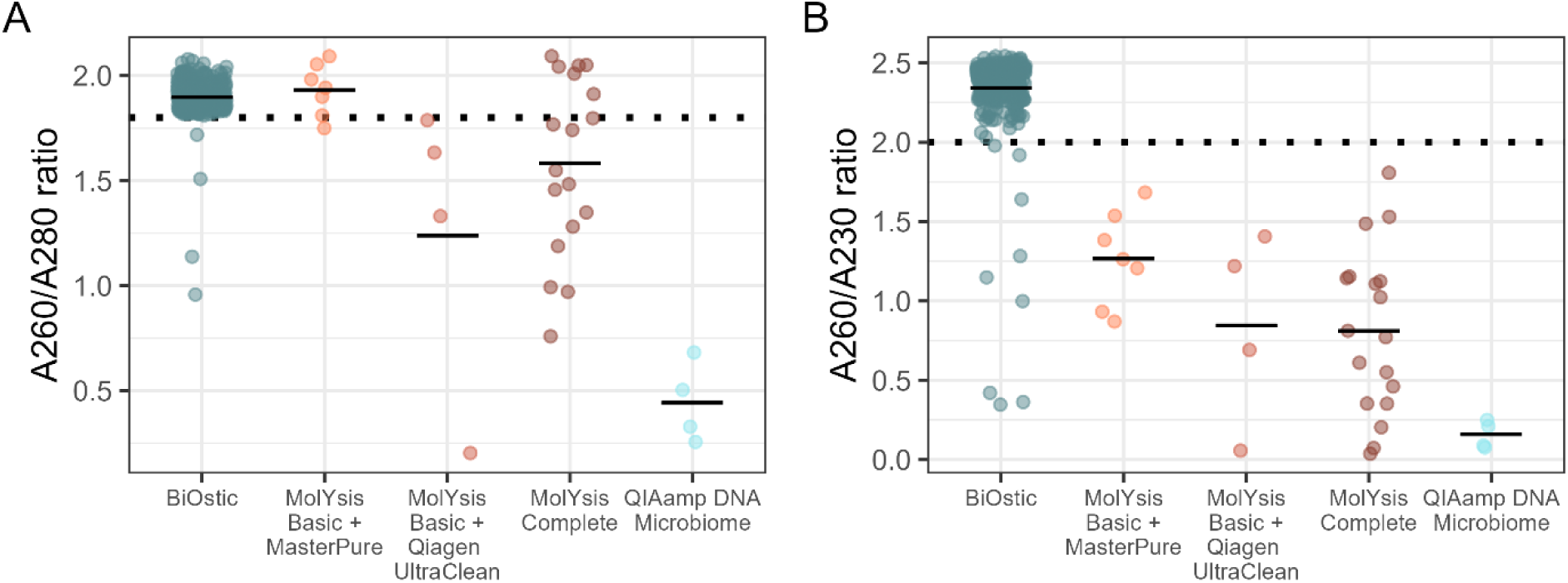
DNA purity across different DNA extraction kits for (**A**) A_260_/A_230_ and (**B**) A_260_/A_280_ ratios. Dotted lines indicate recommended values for pure DNA. Solid lines indicate means for each kit. No sequencing data was obtained with MolYsis extracts nor with QIAamp extracts due to immediate pore death upon library addition to flow cell.

**Supplementary Figure 2.**
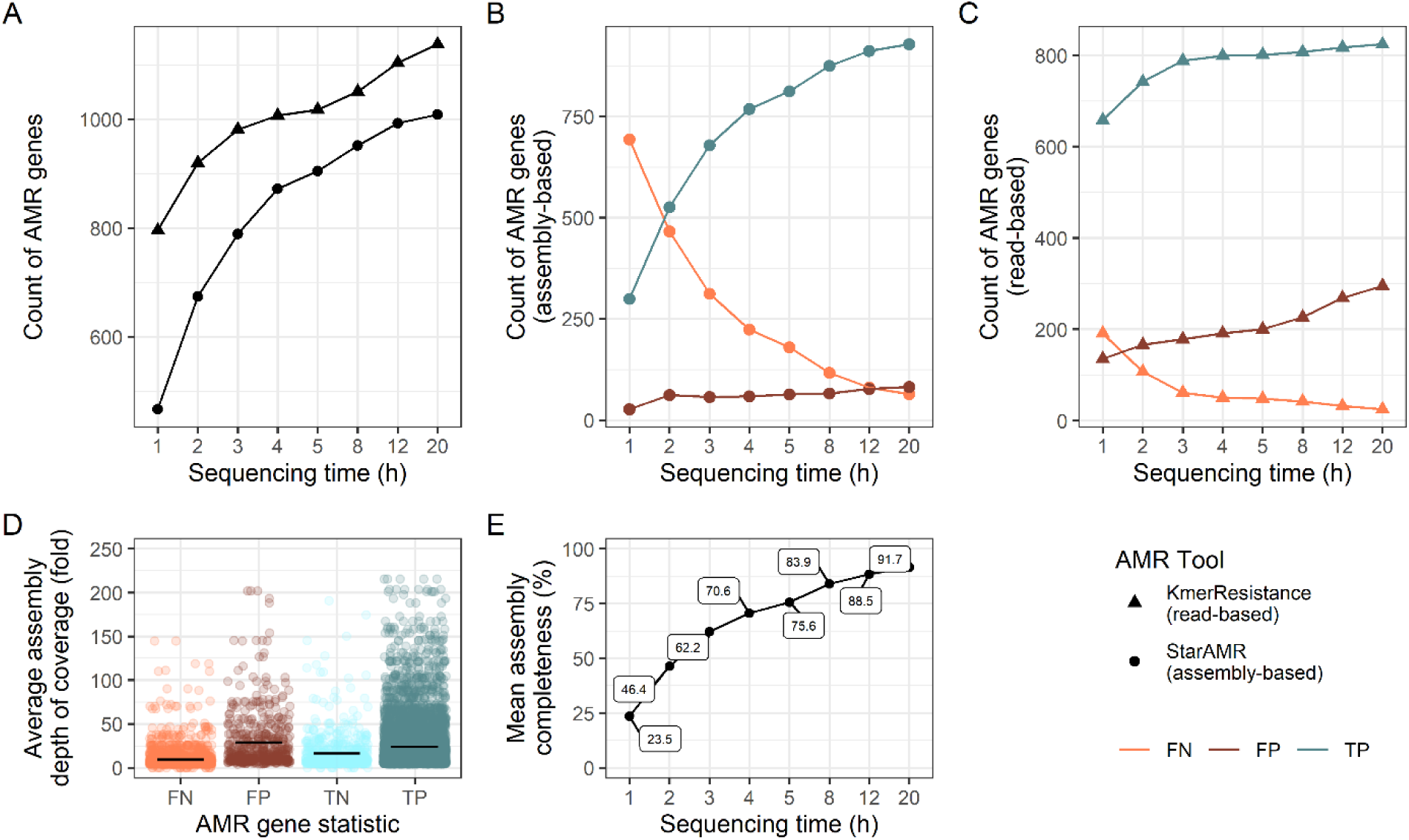
Count of correctly-identified AMR genes in 281 positive BCs using the ONT method. (**A**) Count of total AMR genes detected by both read-based and assembly-based tools. (**B**) Count of AMR genes detected by assembly-based tool (StarAMR) and (**C**) read-based tool (KmerResistance). The reference dataset was matching pure isolates sequenced by Illumina and assemblies analyzed for AMR genes with StarAMR and KmerResistance. The total number of AMR genes detected is 992 in assemblies and 849 in reads, and correctly-identified genes include the correct allelic variant. FosB and pbp5 are not included in read-based KmerResistance database and were excluded, and *bla*_SHV_ alleles were also excluded as many do not have described phenotypes in the ResFinder database but depending on the variant can encode resistance to multiple antimicrobials. (**D**) Average assembly depth of coverage for positive or missed detections of AMR genes. Lines represent the mean. (**E**) Mean assembly completeness over sequencing time, which mirrors assembly-based AMR gene detection. FN = false negative (gene missed in ONT method), TP = true positive (gene correctly detected), FP = false positive (gene detected in ONT method that was not detected in Illumina).

**Supplementary Figure 3.**
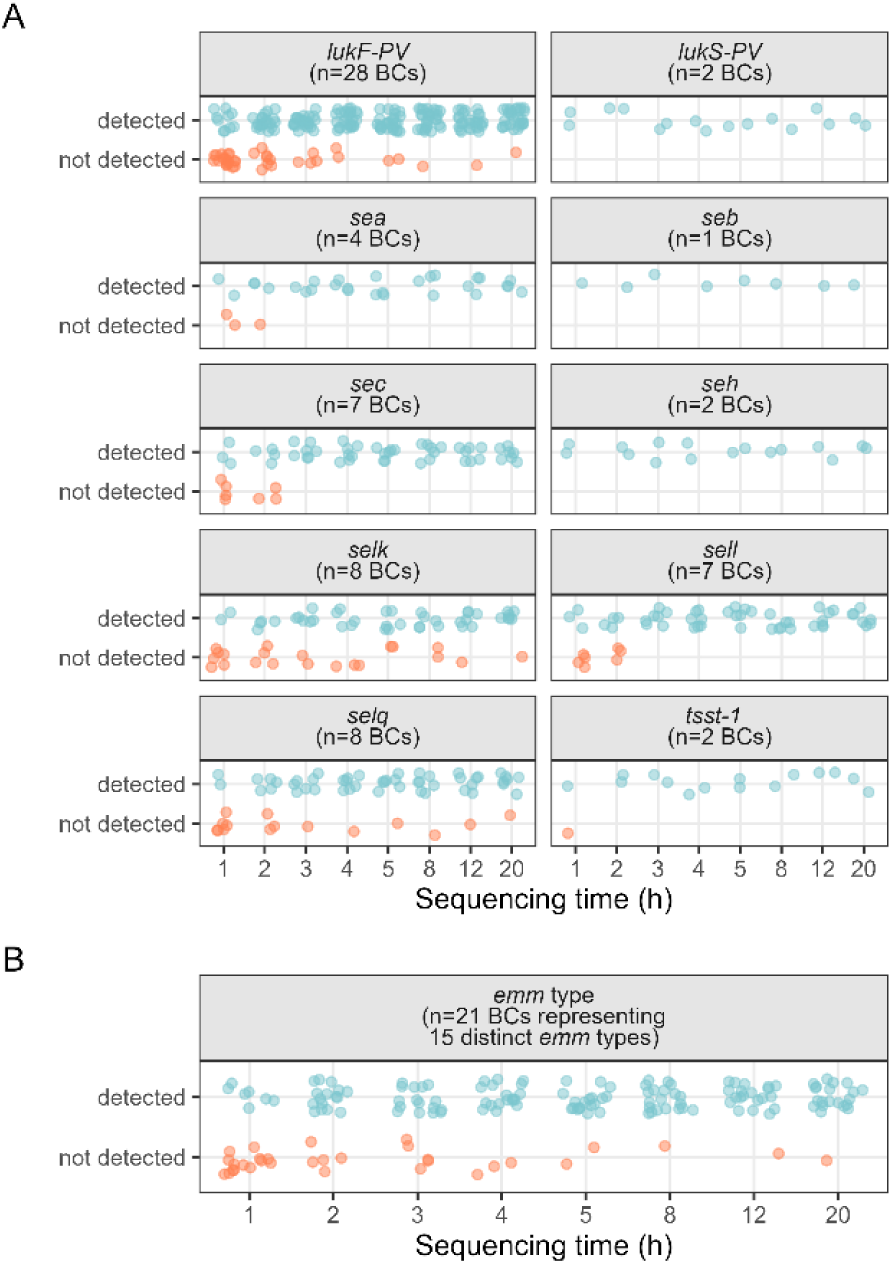
Time to detection for **(A)** *S. aureus* toxin genes across 30 positive blood cultures (BCs) and (**B**) *S. pyogenes emm* type across 21 BCs. Each point represents an individual BC.

**Supplementary Figure 4.**
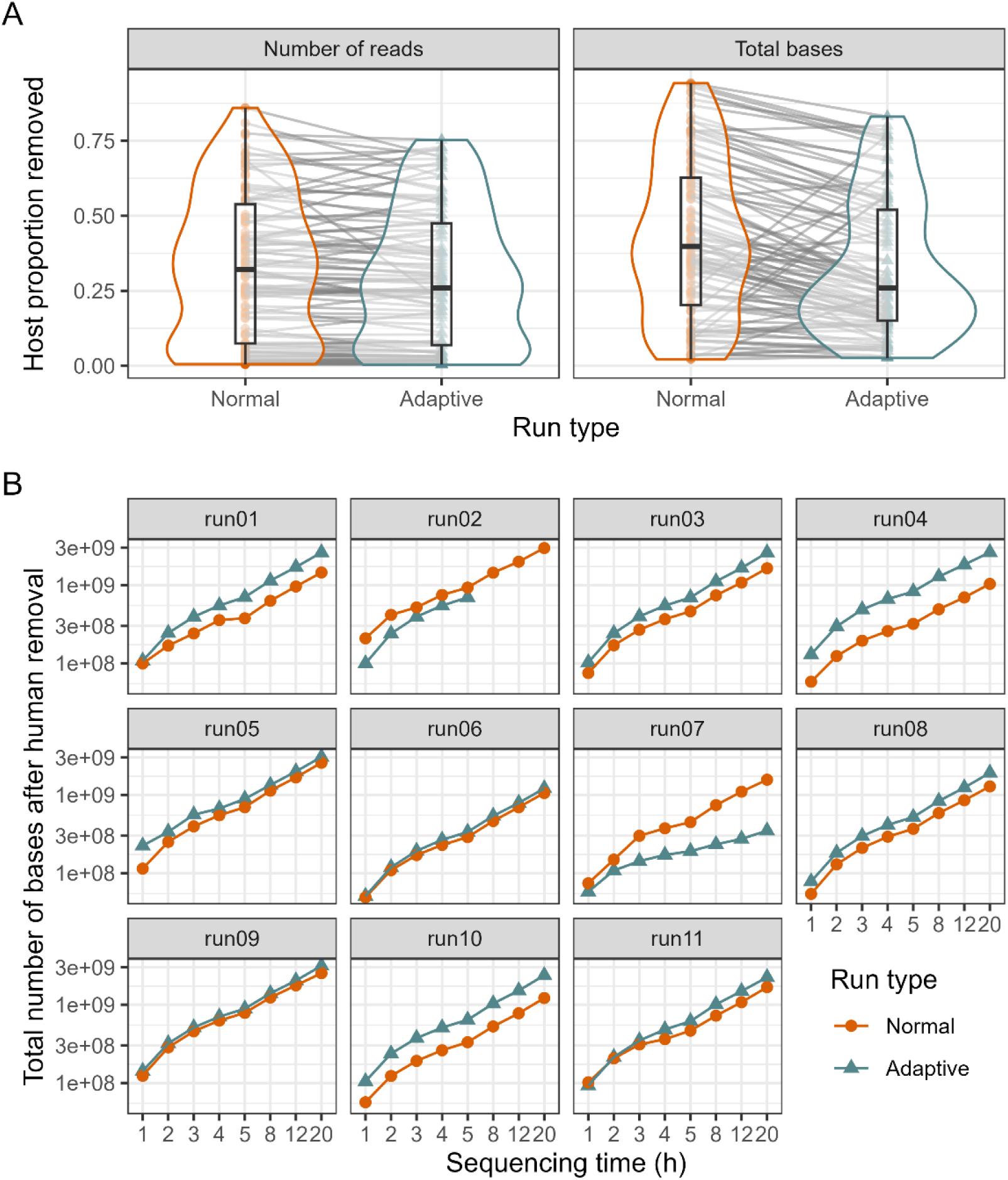
(**A**) Number of host reads (left panel) and bases (right panel) removed during analysis in adaptive vs normal (non-adaptive) runs. Lines represent 25^th^, median, and 75^th^ percentiles. (**B**) Average number of microbial bases sequenced over time (post host depletion) faceted by adaptive run. Each point represents the mean of 4 – 12 BCs.

**Supplementary Figure 5.**
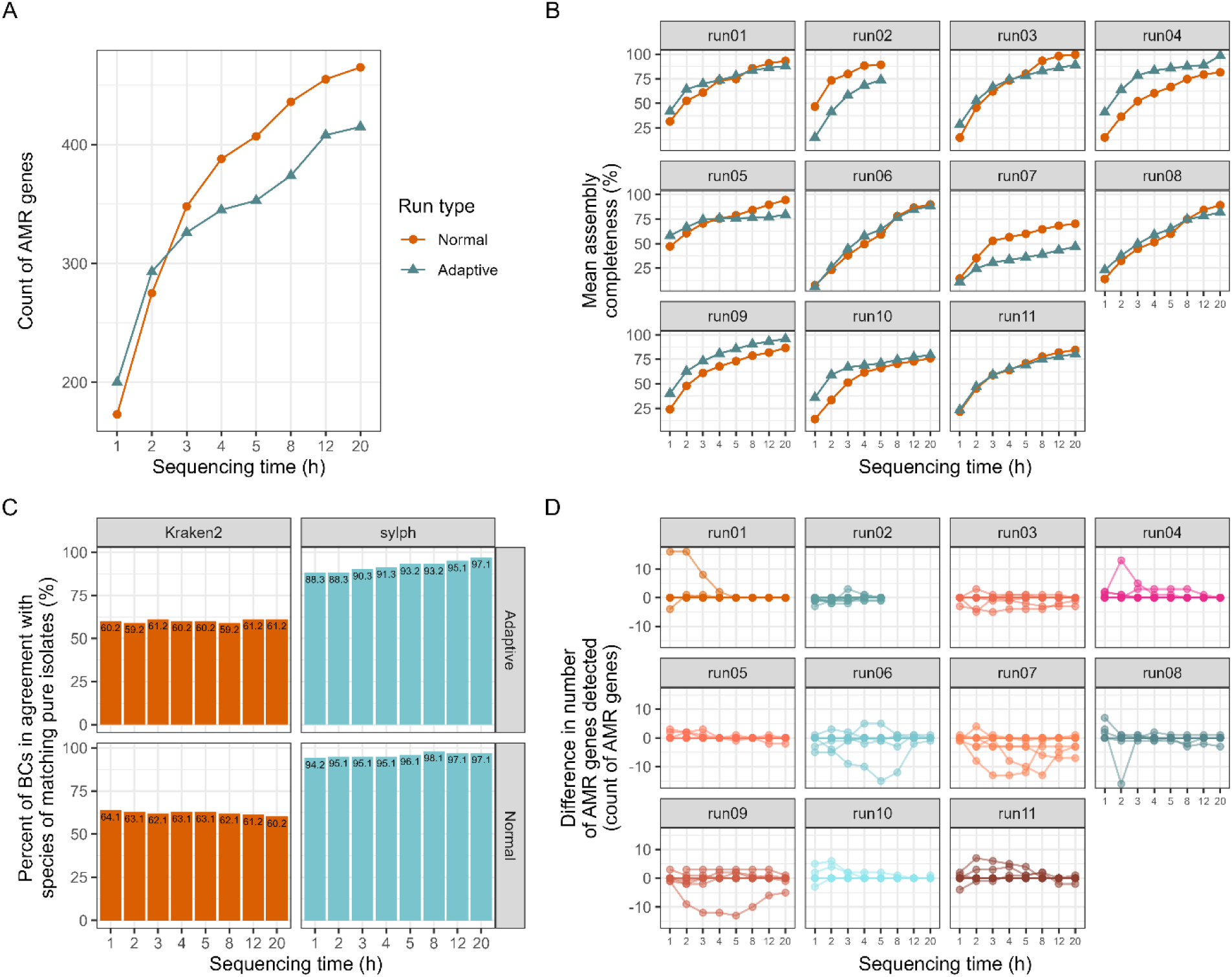
Data output by adaptive sampling runs compared to non-adaptive runs. “Normal” indicates non-adaptive sampling whereas “Adaptive” indicates the libraries were sequenced with host depletion. (**A**) The total number of AMR genes detected over time by assembly-based tool (StarAMR) across all runs for normal and adaptive runs. (**B**) Difference in the number of AMR genes detected across all samples in each run, faceted by each run (n=11). Values above 0 indicate more AMR genes in adaptive data and values below 0 indicate more AMR genes in non-adaptive/normal data. (**C**) Proportion of BCs in agreement with genus (top panel) and species (bottom panel) identity of matching pure isolates over time using Kraken2 (left panel) and sylph (right panel). (**D**) Mean assembly completeness over sequencing time faceted by run. Each run contained between 4 – 12 BCs.

